# Detection and genomic characterisation of a novel hantavirus in Australian dolphins

**DOI:** 10.64898/2026.06.03.729993

**Authors:** Kate Van Brussel, Erin Harvey, Josephine Rieken, Hannah Bender, Jane Hall, Heather Fenton, Karrie Rose, Edward C Holmes

**Author notes:** Author for Correspondence: Prof. Edward C. Holmes, School of Medical Sciences, University of Sydney, Sydney, NSW 2006. Australia.

## Abstract

We report the detection of a novel hantavirus in the lung tissue of two diseased Australian dolphins with histopathological changes. Phylogenetic analysis placed this virus within the genus *Mobatvirus*. This highlights the ability of hantaviruses to infect non-terrestrial mammals and the potential role of marine mammals as one health sentinels.

---

The *Hantaviridae* family of negative-sense RNA viruses comprises four subfamilies and eight genera that have been detected in mammals, fish and reptiles. The genus *Orthohantavirus* (subfamily *Mammantavirinae*) contains the only known zoonotic hantaviruses that occasionally spread to humans via small mammals. Infections in humans are associated with two disease syndromes depending on the virus species and its geographic distribution: haemorrhagic fever with renal syndrome (Europe and Asia) and hantavirus pulmonary syndrome (Americas), which has a high case fatality rate. Far less is known about other genera within the *Mammantavirinae*.

Hantaviruses of the genus *Mobatvirus* have been detected in bat species from Asia, as well as moles and shrews in Europe (*1–5*). Robina virus (*Mobatvirus robinaense*) was first detected in the brain tissue of a black flying fox in Queensland, Australia (*6*). While this animal presented with neurological signs and mild encephalitis, the role of Robina virus in that pathology is unclear (*6*). To date, hantaviruses have not been identified in Australian rodents although antibodies have been detected (*7*). Similarly, hantaviruses are yet to be reported in marine mammals globally. Here, we report the detection of a novel hantavirus in the tissue of two Australian dolphins.

## The Study

### Case 1 (TARZ-13870.1)

The carcass of a young adult male common dolphin (*Delphinus delphis*) was found on a beach in Forster, New South Wales, Australia in December 2020. The animal appeared underweight with below average blubber depths, a mildly ulcerated tongue and lungs that failed to collapse with obvious rib impressions (Figure 1). Histopathology revealed significant changes in the heart characterised by cardiomyocyte degeneration and interstitial infiltration of lymphocytes and plasma cells (Figure 1). Mild, chronic lymphoplasmacytic inflammation was observed in the pleura, tongue, muscularis layer of the colon, and skin associated with degenerative changes and inflammation. Pancreatic trematode infection was also detected associated with per-ductular lymphoplasmacytic cell inflammation that was interpreted as an incidental finding. Additional ancillary testing did not detect *Coxiella burnetii*, cetacean morbillivirus, Influenza A virus, or *Brucella sp*. by PCR (Supplementary Material).

**Figure 1.**
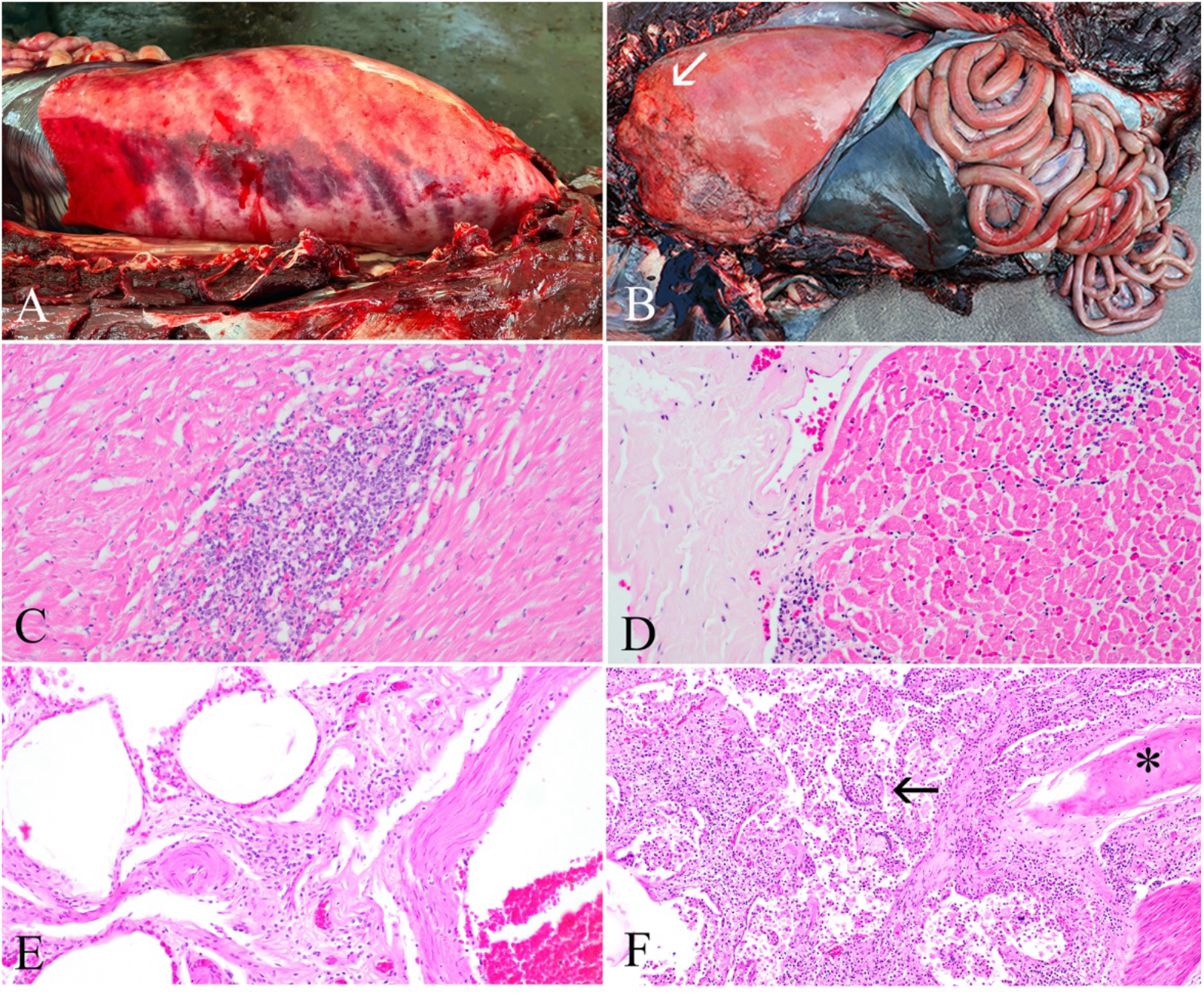
Gross images (A, B) of the thoracic cavity and histopathology (C-F) of the heart and lung of an adult male common dolphin (TARZ-13870.1) and Indo-Pacific bottlenose dolphin (TARZ-16552.1). All histopathology slides (C-F) are at 20x objective and stained with hematoxylin and eosin. (A) Opened thoracic cavity from TARZ-13870.1 and (B) from TARZ-16552.1 with a region of ongoing inflammation of the heart and lung highlighted by the white arrow. (C) Microscopic section of heart from TARZ-13870.1 showing focal aggregates of lymphocytes and plasma cells associated with vacuolation and disruption of cardiomyocytes. (D) Microscopic section of heart from TARZ-16552.1 showing the peri and epicardial fibrosis on the surface of the heart as well as multifocal lymphoplasmacytic myocarditis associated with mild vacuolation of cardiomyocytes. (E) Microscopic section of lung from TARZ-13870.1 showing mild interstitial fibrosis and lymphocytes and plasma cell infiltrates. (F) Microscopic section of lung from TARZ-16552.1 showing multifocal to coalescing neutrophilic, eosinophilic and granulomatous cellular infiltrate amongst sloughed cells, eosinophilic fluid and multiple larval filarid nematodes (black arrow) within an air space. Cartilaginous portion of bronchiole indicated with an asterisk (terminal airways are supported by cartilage in cetaceans). The interstitium is also thickened with fibrosis and lymphocytes and plasma cell infiltrates.

### Case 2 (TARZ-16552.1)

An adult male Indo-Pacific bottlenose dolphin (*Tursiops aduncus*) was found dead on a beach in Kurnell, New South Wales, Australia in 2025. The gross necropsy findings in this individual included poor nutritional condition, advanced tooth wear and evidence of several concurrent infections associated with a compromised immune system, including intravascular filariid nematodes, granulomatous pneumonia and severe chronic fungal dermatitis. Possible intracytoplasmic inclusion bodies associated with epithelial hyperplasia and vacuolar degenerative changes were noted in the tongue. Additional microscopic changes included chronic, mild lymphoplasmacytic myocarditis, interstitial nephritis, gastritis, and meningitis, as well as chronic papillary cystitis (Figure 1). Gram-negative rod-shaped bacteria were localised within the pulmonary granulomas, trachea, and lymph node although culture was not performed. Influenza A virus was not detected via PCR (Supplementary Material). Mild burdens of parasites included tracheal nematodes, subcutaneous plerocercoid cestodes and acanthocephalans and cestodes within the proximal small intestine, all of which are considered incidental in free-ranging dolphins.

Tissue samples were collected from both dolphins for metatranscriptomic sequencing (Supplementary Material). Viruses from the following families were identified: *Anelloviridae, Flaviviridae* and *Polyomaviridae* (TARZ-16552.1), and *Hantaviridae* (TARZ-13870.1 and TARZ-16552.1). Herein, we focus on the hantaviruses. For both cases, the full sequence of the S (nucleocapsid), M (glycoprotein) and L (RNA-dependent RNA polymerase) segments of a novel hantavirus was obtained from the lung metatranscriptome. The number of mapped reads were 2,756 for the S, 16,718 for the M and 3,528 for the L segments in the TARZ-13870.1 lung library, constituting 0.001%, 0.009% and 0.002% of the total library reads respectively. RT-PCR confirmed hantavirus RNA in the spleen, kidney and brain, although the liver, tongue and lymph node were negative (Supplementary Material). No heart tissue was available for this individual. The TARZ-16552.1 lung library mapped read counts were 2,446 for the S, 4,488 for the M and 7,869 for the L segments and 0.001%, 0.003% and 0.004% respectively. RT-PCR of the heart, spleen and lymph node were negative for TARZ-16552.1.

The raw read data for the samples generated are present on the Sequence Read Archive (SRA) under BioProject PRJNA1474109 while the virus consensus sequences are assigned NCBI GenBank accession numbers PZ492986 - PZ492991.

The nucleocapsid sequence of this novel hantavirus shared 66% amino acid identity to Quezon virus (*Mobatvirus quezonense*), identified in Geoffroy’s rousette bat (*Rousettus amplexicaudatus*) in the Philippines (3), and 54% and 71% to Robina virus over the glycoprotein and RNA-dependent RNA polymerase, respectively. Within the phylogeny these viruses formed a distinct clade in the genus *Mobatvirus* (Figure 2). The hantavirus genomes from both cases share 79% nucleotide and 95% amino acid identity over the concatenated genes, suggesting they are variants of the same species provisionally named “delphi hantavirus”.

**Figure 2.**
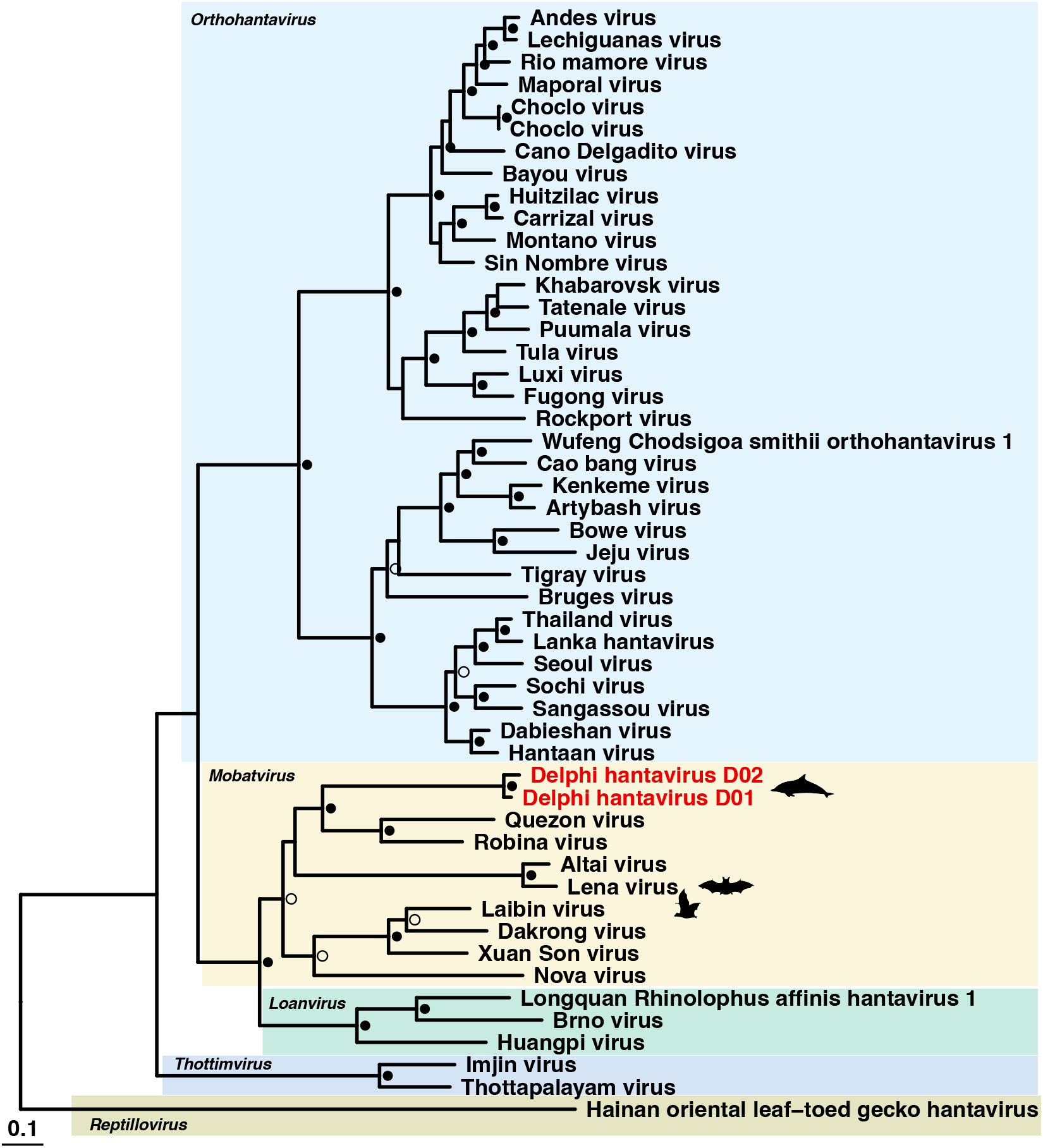
Phylogenetic relationships of the *Hantaviridae* (*Mammantavirinae*) inferred using the concatenated nucleocapsid/glycoprotein/RNA-dependent RNA polymerase genes and the maximum likelihood method. The novel dolphin hantavirus is highlighted in red, the tree is rooted at Hainan oriental leaf-toed gecko hantavirus, and the scale bar represents the number of amino acid substitutions per site. Genera are highlighted and satisfactory bootstrap support is represented by the open circle (<80% and >95% or >80% and <95% SH-like approximate likelihood ratio test and Ultrafast bootstrap approximation) and the closed circle (>80% and >95% SH-like approximate likelihood ratio test and Ultrafast bootstrap approximation).

## Conclusions

We report the detection of a hantavirus in two dolphins with evidence of poor health and concurrent infections, although the impact of hantavirus infection was unclear. Reports of pathology in hantavirus infected non-human animals are sparse, although acute fatal neurological disease is reported in newborn and immunodeficient laboratory rodents (*8*). In the absence of *in situ* diagnostic tests, the role of delphi hantavirus in the pathology observed in both dolphins is unknown. The extent of concurrent illness with multiple pathogens detected in each animal in addition to “normal” parasitic fauna makes it impossible to elucidate which infectious agent(s) were responsible for the lesions. Although mild lymphoplasmacytic inflammatory lesions were noted in cardiorespiratory and neurological tissues (Figure 1), they could not be definitively attributed to hantavirus. The histological lesions in these dolphins did not resemble the severe acute necrotising lesions and vasculitis reported in affected humans and mice (*8*).

Immunosuppression and multiple concurrent stressors may have contributed to the poor overall health of both animals, as previously documented in free-ranging marine mammals (*9-12*). The detection of delphi hantavirus RNA in multiple tissues from TARZ-13870.1 suggests that infection may have been systemic in this individual.

Hantavirus transmission to humans is commonly associated with exposure to rodent urine and droppings, with suspected transmission via the respiratory route. In a marine environment, aspiration of virus particles via the blowhole from a contaminated environment seems plausible, although ingestion from contaminated water is also possible. The mechanism of transmission to cetaceans is currently unknown and warrants further investigation.

The detection of this virus in two different dolphin species suggests possible circulation within the marine environment more broadly. Phylogenetic analysis places both delphi hantavirus variants within the *Mobatvirus* genus and indicates the presence of a cetacean lineage within the *Hantaviridae*, highlighting the importance of continued screening of wildlife population for viruses of zoonotic potential. As marine mammals are key sentinels of ocean health and are facing multiple concurrent stressors, it is likely that emergent pathogens will continue to be identified in these hosts, particularly in degraded environments.

This study highlights the importance of passive wildlife health surveillance programs for early detection of emerging and potentially zoonotic diseases. This surveillance is increasingly valuable in the face of projected anthropogenic impacts, such as increased run-off contributing to pathogen and toxicological pollution and climate change. Monitoring of the evolution and genetic diversity of viruses, particularly at species interfaces, is also essential for pandemic preparedness and public health responses.

## Acknowledgements

The Australian Registry of Wildlife Health is a 40-year science program of Taronga Conservation Society which contributes to the surveillance marine species health in partnership with the New South Wales Departments of Primary Industries and Regional Development and Climate Change, Energy, the Environment and Water. We thank the previous and current staff and volunteers in assisting with managing samples and data related to wildlife health programs, particularly Drs. Peter Johnson and Bryn Lynar, and Nicole Dobson, Allison Rubie, and Karina Hammond. We acknowledge the University of Sydney Veterinary Pathology Diagnostic Services for their assistance in processing the histopathology slides, and researchers at the Australian Centre for Disease Preparedness and Elizabeth Macarthur Agricultural Institute for performing pathogen screening. This work was funded by a National Health & Medical Research Council (Australia) Investigator grant (GNT2017197) and an Australian Research Council Discovery Project grant (DP240101313) awarded to ECH. Wildlife Health Australia provided funding assistance through their One Health Fund for ancillary testing on TARZ-13870.1.

