## Supplementary Information for "Detection and genomic characterisation of a novel hantavirus in Australian dolphins"

### Sample collection and histopathology

Gross necropsies and histopathology were performed by the Australian Registry of Wildlife Health. Fresh tissues were collected and placed at -80°C within three hours of post-mortem examination then transported to the University of Sydney for RNA extraction and library processing. Additionally, representative tissue samples were taken in 10% neutral buffered formalin, routinely processed into paraffin-embedded blocks, sectioned into 5-micron sections and stained with hematoxylin and eosin, as well as select special stains (Gram, Periodic Acid Schiff, and Ziehl-Neelsen).

### Sample and library processing

Total RNA was extracted from tissue samples using the RNeasy Plus Mini Kit (Qiagen) and sequencing libraries were generated using the Illumina Stranded Total RNA Prep with Ribo-Zero Plus rRNA depletion at the Australian Genome Research Facility, Melbourne. Libraries for TARZ-13870.1 were sequenced on the NextSeq2000R as 100bp paired-end reads and for TARZ-16552.1 on the NovaSeqX Plus as 150bp paired-end. Reads were quality trimmed using Cutadapt (version 5.0), *de novo* assembled into contigs using MEGAHIT (version 1.2.9) and compared against the Reference Viral Database and nr database using DIAMOND BLASTX (version 2.1.10) (*1–5*). Mapped read counts were calculated from the trimmed reads using RSEM (version 1.3.3) (*6*). Raw reads were deposited onto the NCBI Sequence Read Archive database under BioProject number PRJNA1474109, BioSample numbers SAMN60586720 to SAMN60586724.

### Phylogenetic analysis

The nucleocapsid, glycoprotein and RNA-dependent RNA polymerase genes from 48 hantaviruses sequences were downloaded from NCBI GenBank. These sequences and delphi hantavirus were concatenated into a nucleocapsid/glycoprotein/RNA-dependent RNA polymerase amino acid sequence and aligned using the E-INS-I algorithm within MAFFT (version 7.490) (*7*). Maximum likelihood trees were inferred from these data using IQ-Tree (version 2025.1.2), employing Model Finder to find the best-fit model of amino acid substitution (*8,9*). Nodal support was calculated using the SH-like approximate likelihood ratio test and Ultrafast bootstrap approximation (1000 replicates) (*8,10*). Nearest neighbour interchange (NNI) was used to control for overestimation of Ultrafast bootstrap values (*8*).

### RT-PCR of additional tissues

The SuperScript IV One-Step RT-PCR System (Invitrogen) was used to perform RT-PCR on the total RNA. The primers DH01F – 5' CTTAAAGAGAGAGCATCC 3' and DH01R – 5' CCAAATGTTAGGACATAATC 3' targeting the nucleocapsid of delphi hantavirus D01 was used to screen the brain, spleen, liver, lymph node, tongue and kidney of TARZ-13870.1. The primers DH02F – 5' AGGTCCTTACTAAATTTTCTGAGA 3' and DH02R – 5' TTAGTCCTAGCAGTCTAACCTT 3' targeting the RNA-dependent RNA polymerase of delphi hantavirus D02 was used to screen the lung, heart, spleen and lymph node of TARZ-16552.1. The annealing temperature and extension time for both primers pairs were 60$^{o}$C and 30 seconds, respectively.

Viral RNA was extracted from tissue samples using the MagMax 96 Viral RNA Isolation Kit (Thremo Fisher Scientific) and the SuperScript III One-Step RT-PCR System (Invitrogen). The viral RNA was used for screening of Influenza virus A, *Brucella sp.* and morbillivirus at the Australian Centre for Disease Preparedness, as described previously (*11*). *Coxiella* *burnetti* qPCR was performed at the Elizabeth Macarthur Agricultural institute using previously published primers and cycling conditions (*12–14*).
